# Cytotoxic Activity of CD4 T Cells During the Early Stage of Autoimmune Neuroinflammation

**DOI:** 10.1101/2020.03.10.985614

**Authors:** Fernando Pradella, Vinicius O. Boldrini, Ana Maria Marques, Guilherme A. D. Morais, Carolina Francelin, Rani S. Cocenza, Vitor C. Lima, Maurilio Bonora, Natalia S. Brunetti, Bruna B. Campos, Evelise S. M. Fonseca, Michelle Rocha-Parise, Carla R. V. Stella, Alfredo Damasceno, Felipe von Glehn, Ana Leda F. Longhini, Leonilda M. B. Santos, Alessandro S. Farias

## Abstract

Pathogenic CD4+ T cells are capable of initiating neuroinflammation in experimental autoimmune encephalomyelitis (EAE). However, the precise effector mechanism of these autoaggressive CD4+ T cells is not entirely elucidated. Here, we demonstrated that pathogenic CD4+ T cells, upon autoantigen stimulation, developed a cytotoxic phenotype at the onset of EAE. The cytotoxic activity of pathogenic CD4+ T cells was sufficient to explain the initial myelin lesion. Consistently, CD4+ T cells of peripheral blood (PBMCs) and cerebrospinal fluid (CSF) from relapse-remitting multiple sclerosis (RRMS) patients present an enhancement of the cytotoxic profile in comparison with healthy control (HC). Moreover, cytotoxic CD4+ T cells (CD4-CTLs) are restrained in the PBMCs of Natalizumab-treated RRMS patients. Mechanistically, autoaggressive CD4-CTLs matched the majority of the molecular pathways of effector CD8+ T cells. Altogether, our findings point to potential new targets for monitoring MS diagnosis, treatment, and the development of novel therapeutic avenues.

## Introduction

Experimental autoimmune encephalomyelitis (EAE) is a widely accepted animal model of multiple sclerosis (MS). EAE shares many pathophysiological features with MS, such as chronic neuroinflammation, demyelination, neuronal damage, and is generated by the autoimmune attack on the central nervous system (CNS) (Baxter, 2007; Steinman and Zamvil, 2006). The disease can be induced in susceptible animals by active immunization with self-epitopes of myelin or by the passive transfer of pre-activated lymphocytes (Paterson, 1960; Rivers et al., 1933). More specifically, EAE can be induced by the adoptive transfer of CD4+ T cells specific to the neuroantigen (Ben-Nun et al., 1981; Flugel et al., 1999; Schluesener and Wekerle, 1985). Even though the inflammatory activity of these cells is well known, the mechanisms by which encephalitogenic CD4+ T cells initiate tissue lesions are not entirely elucidated (Glatigny and Bettelli, 2018; Wagner et al., 2019). It is not clear if these cells are capable of promoting direct cytolytic activity over the target tissue or if they affect the target tissue by creating an inflammatory milieu allowing other leucocytes entry in the CNS to exert their effector function.

Although CD8+ T cells do not imply the onset of EAE (Jiang et al., 1992; Koh et al., 1992; Saligrama et al., 2019), the blockage or deletion of cytotoxic-related molecules, especially transcription factors, results in a minor or delayed onset of the disease (Aqel et al., 2018; Martinez Gomez et al., 2012; Nohara et al., 2001; Y. Wang et al., 2014). Although it is not particularly new (Irle et al., 1984; Ozdemirli et al., 1992; Wagner et al., 1975; 1977), a growing body of evidence supports the cytotoxic activity of CD4+ T cells during chronic viral infections, tumor-specific immune response, and allograft rejection (Caielli et al., 2019; Fleischhauer et al., 2001; Quezada et al., 2010; Serroukh et al., 2018; Śledzińska et al., 2020; Wilkinson et al., 2012).

In 2007, Kebir and colleagues showed that the cytotoxic activity of encephalitogenic CD4+ T cells in the EAE model is essential for breaching the blood-brain barrier (Kebir et al., 2007). Furthermore, treating EAE with Serpina3n, a granzyme B (GzmB)-inhibitor, resulted in a significant reduction of EAE severity, although infiltration of T cells into the CNS was not reduced (Haile et al., 2015). Recent studies indicate a role of cytotoxic CD4+ T cells (CD4-CTLs) during the course of RRMS as well (Beltrán et al., 2019; Kebir et al., 2007; Peeters et al., 2017; Schafflick et al., 2020; Zaguia et al., 2013).

In this context, we investigated the role of CD4-CTLs during the early phase of autoimmune neuroinflammation. We reported that during the early stage of EAE encephalitogenic CD4+ T cells build a cytotoxic profile, which is enhanced after these cells reach the CNS. These encephalitogenic CD4-CTLs alone were capable of initiating CNS damage in a healthy CNS tissue. Moreover, our results in the experimental model were consistently replicated in CD4+ T cells of peripheral blood (PBMCs) and cerebrospinal fluid (CSF) from RRMS patients. The enhancement of CD4-CTLs presented potential implications for the diagnosis and treatment of RRMS.

## Results

### Enhancement of the cytotoxic activity of encephalitogenic CD4+ T during EAE

In order to evaluate the cytotoxic profile of encephalitogenic CD4+ T cells, we sorted CD3+CD4+ cells from lymph nodes (LN) and central nervous system (CNS) of mice immunized with neuroantigen (MOG_35-55_), or with non-self-antigen [ovalbumin (OVA)]. Then, we analyzed the expression of classical proinflammatory and cytotoxic-related molecules by qPCR. In general, CD4+ T cells sorted from MOG_35-55_-immunized mice presented a significantly higher expression of cytotoxic-related molecules in comparison with OVA-immunized animals either on the LN or CNS (**Fig. 1A**).

**Figure 1.**
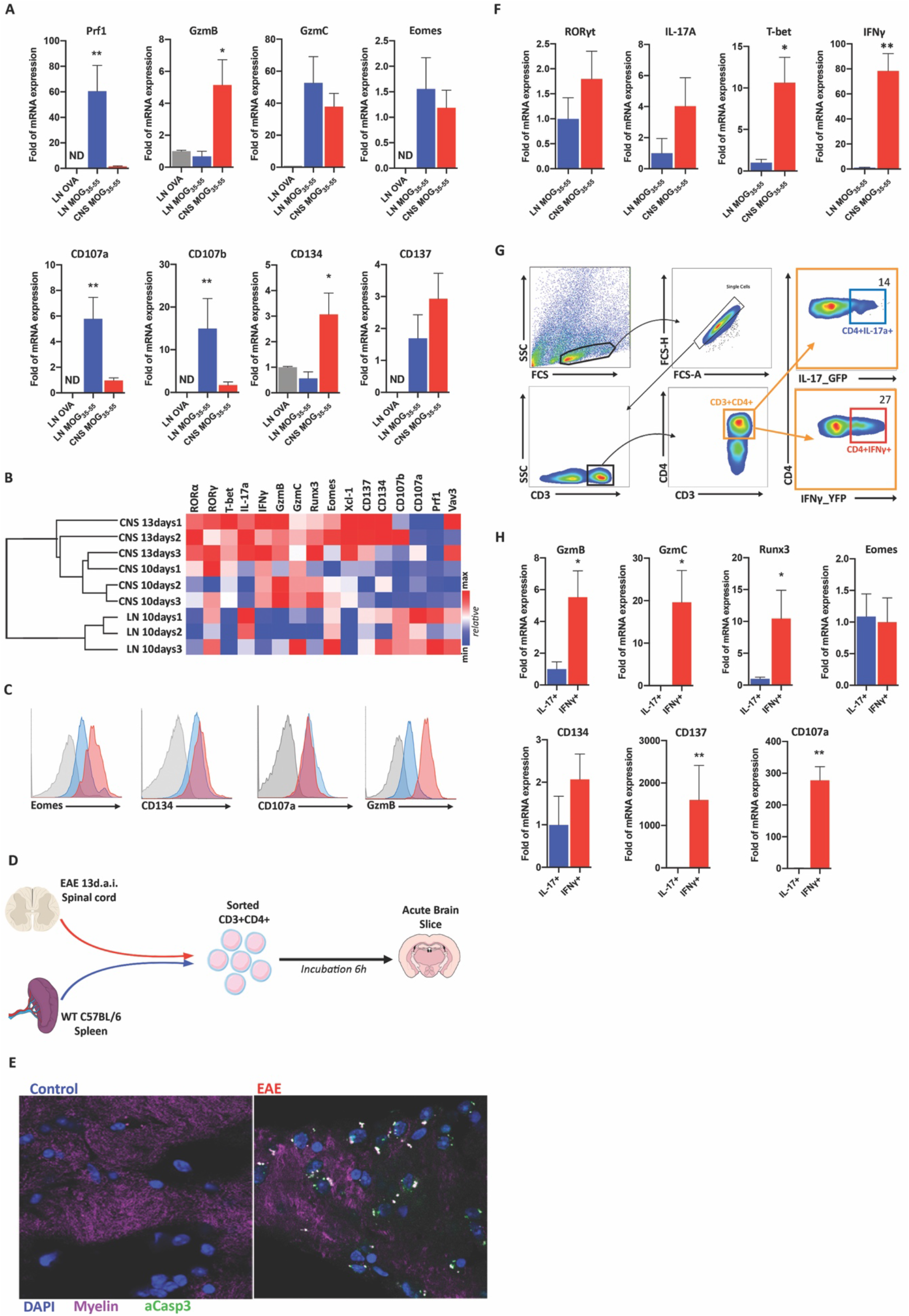
Cytotoxic activity of CD4+ T cells during the clinical evolution of EAE. **(A)** mRNA expression of cytotoxic-related molecules in CD4+ cells sorted from the lymph nodes (LN) of OVA-immunized 10 d.a.i. (gray), LN of MOG_35-55_-Immunized mice; 10 d.a.i. (Blue) or CNS of MOG_35-55_-Immunized mice; 10-13 d.a.i. (red) (*n=3-9*). **(B)** Hierarchic cluster analysis of cytotoxic (Eomes, Runx3, GzmB, GzmC, Prf1, CD107a, CD107b, CD134, CD137 and VAV3) and inflammation-related molecules (T-bet, RORα, RORγt, IFNγ, IL-17a and XCL-1) in sorted CD4+ T cells during the evolution (lymph nodes 10 d.a.i. (LN10days), spinal cord 10 d.a.i. (CNS10days) and spinal cord 13 d.a.i. (CNS13days)) of EAE. **(C)** Representative flow cytometry analysis of Eomes, CD134, CD107a and GzmB of CD4+ T cells from CNS (red) at 10-13 d.a.i. and lymph nodes (blue) at 10 d.a.i. of EAE and isotype control (gray) (*n=5*) **(D)** Experimental design of acute slice incubation. **(E)** Confocal microscopy (63x) of a brain slice after incubation with CD4+ T cells from the spleen of unimmunized animals (left panel) or CD4+ cells sorted from the spinal cord 13 d.a.i. (right panel) (*n=9*). **(F)** mRNA expression of T-bet, IFNγ, RORγt, and IL-17A of CD4+ T cells sorted LN of MOG_35-55_-Immunized mice; 10 d.a.i. (Blue) or CNS of MOG_35-55_-Immunized mice; 10-13 d.a.i. (red) (*n=3-9*). **(G)** Gate strategy for sorting CD3+CD4+ (orange gate) cells expressing IL-17a (blue gate) or IFNγ (red gate). **(H)** mRNA expression of cytotoxic-related molecules among CD4+IL-17a+ (blue) or CD4+IFNγ+ (red) cells from CNS 10-13 d.a.i. (*n=4-7*). Data are represented in mean +/− SEM; *p<0.05, **p<0.01

Interestingly, cluster analysis demonstrated that CNS infiltrating CD4+ T cells present an enhancement of the cytotoxic profile (**Fig. 1B**). These data were corroborated by flow cytometry (**Fig. 1C**). Nevertheless, the expression of the majority of cytotoxic-related molecules by CD4+ T cells sorted from LN of mice immunized with MOG_35-55_ is significantly higher in comparison with CD4+ T cells sorted from LN of mice immunized with OVA (**Fig. S1A**). These results revealed that CD4+ T cells built a cytotoxic profile during the evolution of EAE upon autoantigen stimulation, which is enhanced after the cells reach the CNS.

Although it is clear that these cells exhibit a cytotoxic profile during the evolution of the disease, this does not necessarily mean that they have cytotoxic activity in the target organ. Thus, intending to evaluate the cytotoxic effector activity of encephalitogenic CD4+ T cells, we incubated acute brain slices with sorted CD3+CD4+ T cells from the CNS of MOG_35-55_-immunized mice 13 days after immunization (d.a.i.), or CD3+CD4+ T cells from the spleen of unimmunized animals (controls) and evaluated the presence of activated caspase-3, which is directly cleaved by GzmB (Goping et al., 2003) (**Fig. 1D**). Our results demonstrated that encephalitogenic CD4+ T cells alone are capable of promoting a direct cytotoxic activity over CNS tissue (**Fig. 1E**).

### Cytokine profile of cytotoxic CD4+ T cells after reach the CNS

Next, we explored the mechanisms by which these cells become cytotoxic and how important they are for the development of the autoaggressive response. Some studies have shown that CD4-CTLs is associated with a Th1 profile (Ozdemirli et al., 1992; Serroukh et al., 2018; Sun and Wekerle, 1986). In contrast, Kebir and colleagues demonstrated that CD4-CTLs present a Th17 profile in the EAE model (Kebir et al., 2007). Our qPCR analysis showed an increased expression of T-bet and IFNγ by CD4+ T cells after reach the CNS (**Fig. 1F**). No significant difference was found in the expression of RORγt of IL-17A (**Fig. 1F**). Then, we used IL-17A^YFP^ or IFNγ^GFP^ reporter mice to evaluate the expression levels of cytotoxic-related molecules in sorted CD4+IL-17a+ or CD4+IFNγ+ T cells after these cells reached the CNS in EAE model (**Fig. 1G**). The expression of the majority of the cytotoxic-related molecules was significantly higher in CD4+ IFNγ-producing T cells about CD4+ IL-17A-producing cells (**Fig. 1H**).

Thus, our data agree with the majority of the literature, which demonstrated a prevalence of Th1-like profile by CD4-CTLs. Nevertheless, Th17 encephalitogenic cells turn to IFNγ-producing cells after entry into the CNS (Hirota et al., 2011; Y. Wang et al., 2014). Therefore, cytotoxic IFNγ-producing CD4+ T cells inside the CNS and cytotoxic Th17 in the periphery might represent the same population in different effector stages. Interestingly, the development of IFNγ-producing Th17 cells required the expression of T-bet and Runx1 or Runx3 (Y. Wang et al., 2014).

### Expression of Runx3 and CD4-CTLs

Runx3 is a crucial transcription factor for T CD8+CD4-lineage differentiation as well as for the initial cytotoxic program inducing the expression of CTL-linage associated genes, including IFNγ (Cheroutre and Husain, 2013; Istaces et al., 2019). Therefore, the enhancement of this transcription factor could explain the cytotoxic activity as well as the Th1-like profile.

Thus, we evaluated the expression of Runx3 by encephalitogenic CD4+ T cells, its relation with the cytotoxic profile, and its importance to the onset of EAE. Our qPCR results demonstrated a significant enhancement of the expression of Runx3 by encephalitogenic CD4+ T cells as well as by IFNγ-producing CD4+ T cells (**Fig. 1H and 2A**). Therefore, we used Runx3 reporter mice (Runx3^YFP^) to evaluate the expression of Runx3 by encephalitogenic CD4+ T cells at the protein level. We observed a significant increase in the expression of Runx3 by CD4+ T cells after reach the CNS (**Fig. 2B**) as well as an enrichment of CD4+Runx3+ T cells infiltrated in the CNS in comparison with CD4+ T cells from LN (**Fig. 2C and 2D**). Also, the expression levels of GzmB was significantly higher in CD4+Runx3+ cells compared to CD4+Runx3-cells infiltrated into the CNS (**Fig. 2E**).

**Figure 2.**
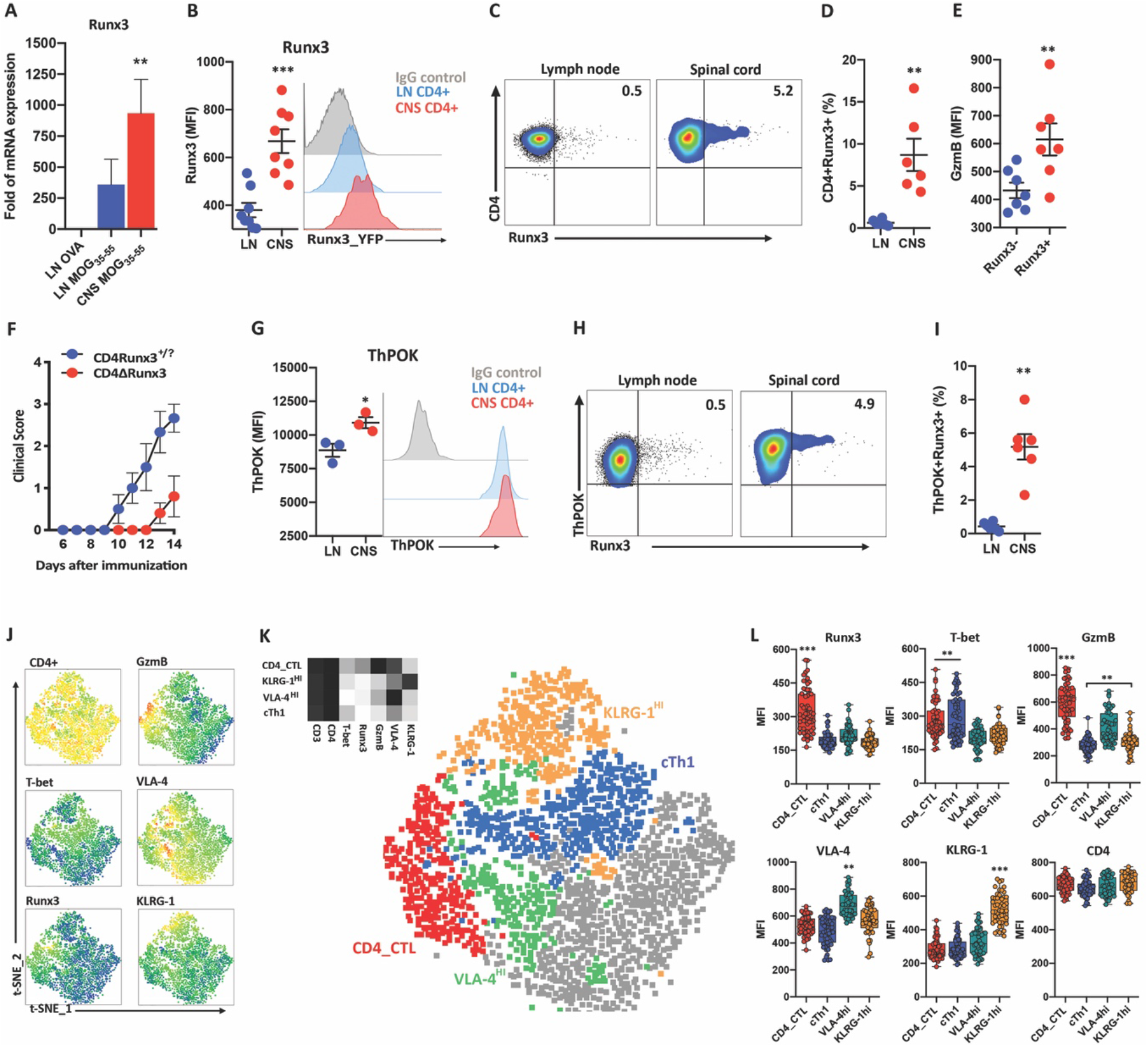
Runx3 expression by CD4+ T cells during the course of EAE. **(A)** mRNA expression of Runx3 in sorted CD3+CD4+ T cells from OVA-immunized 10 d.a.i. (gray), LN of MOG_35-55_-Immunized mice; 10 d.a.i. (Blue) or CNS of MOG_35-55_-Immunized mice; 10-13 d.a.i. (red) (*n=3-9*). **(B)** Runx3 expression (MFI) in CD3+CD4+ cells from LN 10 d.a.i (blue) and CNS 10-13 d.a.i (red). **(C)** Representative flow cytometry analysis of Runx3+ cells among gated CD3+CD4+ cells from LN 10 d.a.i. (left), CNS 10-13 d.a.i. (right) (*representative of n=6*). **(D)** Frequency of Runx3+ cells in gated CD3+CD4+ cells from LN 10 d.a.i. (blue), CNS 10 d.a.i. (red). **(E)** GzmB expression (MFI) in CD3+CD4+Runx3-cells (blue) or CD3+CD4+Runx3+ cells (red). **(F)** Clinical evolution of EAE by CD4Runx3?/+ mice (blue) or CD4∆Runx3 mice (red) immunized with MOG35-55. **(G)** ThPOK expression (MFI) among CD3+CD4+ cells from LN 10 d.a.i (blue) and CNS 10-13 d.a.i (red) – representative of two independent experiments (*n=3 in each experiment*). **(H)** Representative Flow cytometry analysis of Runx3 expression in CD3+CD4+ThPOK+ cells (*n=6*). **(I)** Frequency of Runx3+ cells among gated CD3+CD4+ThPOK+ cells from LN 10d.a.i. (blue), CNS 10d.a.i. (red). **(J)** Heatmap of t-SNE-based overview of CD4, GzmB, T-bet, VLA-4, Runx3 and KLRG-1 expression. (**K**) Mean population expression levels of all markers used for t-SNE visualization and FlowSOM clustering (left). The t-SNE algorithm (10,000 CD3+CD4+ cells, randomly selected from CNS 13 d.a.i. (*n=4*)) was used to depict different 5 populations therein. FlowSOM-based cell populations are overlaid as a color dimension (right). **(L)** Expression intensity (MFI) of Runx3, T-bet, GzmB, VLA-4, KLRG-1 at single cell level (400x CD3+CD4+ cells randomly selected) for each FlowSOM-based population. Data are represented in mean +/− SEM; *p<0.05, **p<0.01 ***p<0.001.

Next, to evaluate the importance of Runx3 by encephalitogenic CD4-CTLs pathogenicity during the onset of EAE, we generated Runx3 tissue-specific (CD4+ cells) knockout mice (CD4∆Runx3). Strikingly, the deletion of Runx3 in CD4+ cells diminished the incidence and severity of EAE (**Fig. 2F**). Moreover, the expression of CD107a by CD4+ T cells in the LN from CD4∆Runx3 mice were significantly decreased (**Fig. S1B**). These data indicate that the expression of Runx3 and the cytotoxic activity of CD4+ T cells contribute to the development of the autoaggressive response in the EAE model, at least during the early stage of the disease. During T cell development in the thymus, the balance between the expression of Runx3 and ThPOK is determinant to CD8 or CD4 lineage fate, respectively. In this context, the expression of Runx3 represses ThPOK and terminates CD4 expression as well (Cheroutre and Husain, 2013; Setoguchi et al., 2008). In contrast, our data showed a slight increase in expression of ThPOK after CD4+ T cell entry in the CNS (**Fig. 2G**) as well as no regulation of CD4 molecule expression (**Fig. 2L**). Besides, we observed that almost all of CD4+Runx3+ T cells were also ThPOK positive (**Fig. 2H and Fig. 2I**) Of note, in the context of human cytomegalovirus infection, Th1 CD4-CTLs have been found to enhance the expression of Runx3 in the absence of ThPOK downregulation as well (Serroukh et al., 2018).

### CD4+ T cells landscape into the CNS

In order to identify the CD4-CTLs infiltrated in the CNS, we used t-distributed stochastic neighbor embedding (t-SNE) to evaluate the co-expression level of CD4, Runx3, T-bet, GzmB, VLA-4 (CD49d, α4β1, α4β7), and KLRG1 in CD3+CD4+ T cells population from CNS during the onset of EAE (**Fig. 2J**). Then, we applied an artificial neural network-based algorithm (FlowSOM) to generate specific populations based on the expression of those markers (**Fig. 2K**). FlowSOM-based nodes were then manually annotated in the CD3+CD4+ t-SNE landscape (**Fig. 2K**). The CD4-CTLs population was defined by the high expression of Runx3, GzmB, and T-bet. These data were confirmed by the expression intensity of those markers at the single-cell level of each annotated population (**Fig. 2L**). Interestingly, the VLA4^hi^ population also present a high expression of GzmB, although it was significantly lower in comparison with CD4-CTLs (**Fig. 2K and Fig. 2L**). Conventional Th1 (cTh1) population was defined by the high expression of T-bet and low expression of Runx3, GzmB, VLA-4, and KLRG1 (**Fig. 2K and Fig. 2L**). We identified one population that expresses high levels of KLRG1 and low levels of Runx3, T-bet, GzmB, and VLA-4 (**Fig. 2K and Fig. 2L**). KLRG1 is highly expressed in short-lived effector CD8+ T cells (Hamilton and Jameson, 2007). Moreover, the expression of KLRG-1 is a direct consequence of T-bet expression (Joshi et al., 2007).

### Cytotoxic profile of CD4+ T cells in PBMCs of the untreated patient with MS

In EAE, the cytotoxic could be detected in the periphery during the early phase of the disease (**Fig. S1**). Therefore, in order to test whether the data generated in the animal model would apply to human disease, we recruited untreated RRMS and matched healthy controls (HC) to evaluate the extension of our data in PBMCs (**Table 1**). Here, we included RRMS patients assessed at the very early stage of the diagnosis (diagnosis) as well as patients that withdrawal the treatment by therapeutic failure or after reach 24 months of Natalizumab (NTZ)-treatment (washout) (**Table 1**). Then, we analyzed the expression of GzmB by CD3+CD4+ or CD3+CD8+ cells in PBMCs by flow cytometry and the expression of cytotoxic-related molecules by qPCR in sorted CD3+CD4+ T cells. Our data demonstrated an increased percentage of CD4+GzmB+ but not CD8+GzmB+ T cells in RRMS patients in relation to HCs (**Fig. 3A - 3D**). Strikingly, there is no correlation between the percentages of CD4+GzmB+ and CD8+GzmB+ T cells, which demonstrates a specific enhancement of the cytotoxic profile by CD4+ T cells (**Fig. 3E)**.

**Table1.**
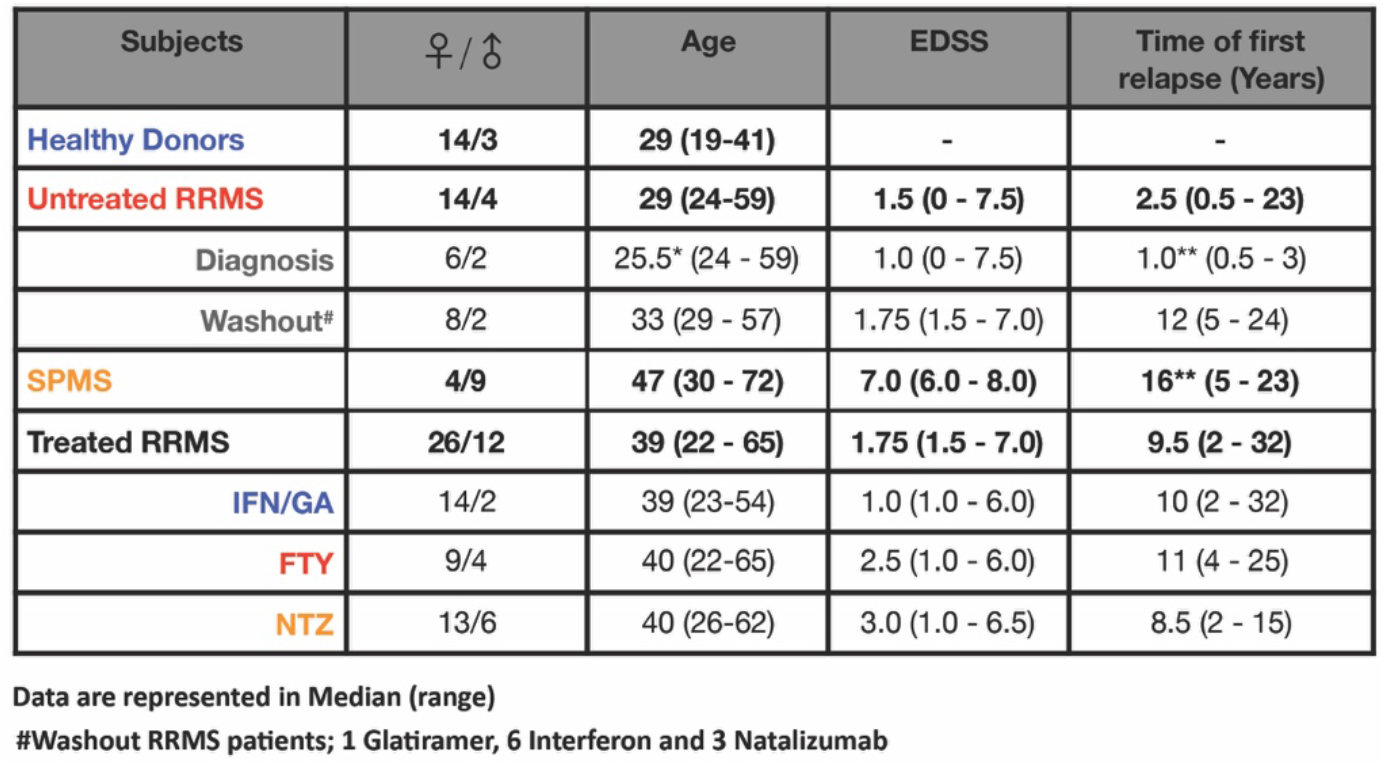

| Subjects | ♀ / ♂ | Age | EDSS | Time of first relapse (Years) |
| --- | --- | --- | --- | --- |
| Healthy Donors | 14/3 | 29 (19-41) | - | - |
| Untreated RRMS | 14/4 | 29 (24-59) | 1.5 (0 - 7.5) | 2.5 (0.5 - 23) |
| Diagnosis | 6/2 | 25.5* (24 - 59) | 1.0 (0 - 7.5) | 1.0** (0.5 - 3) |
| Washout# | 8/2 | 33 (29 - 57) | 1.75 (1.5 - 7.0) | 12 (5 - 24) |
| SPMS | 4/9 | 47 (30 - 72) | 7.0 (6.0 - 8.0) | 16** (5 - 23) |
| Treated RRMS | 26/12 | 39 (22 - 65) | 1.75 (1.5 - 7.0) | 9.5 (2 - 32) |
| IFN/GA | 14/2 | 39 (23-54) | 1.0 (1.0 - 6.0) | 10 (2 - 32) |
| FTY | 9/4 | 40 (22-65) | 2.5 (1.0 - 6.0) | 11 (4 - 25) |
| NTZ | 13/6 | 40 (26-62) | 3.0 (1.0 - 6.5) | 8.5 (2 - 15) |

Data are represented in Median (range)

**Figure 3.**
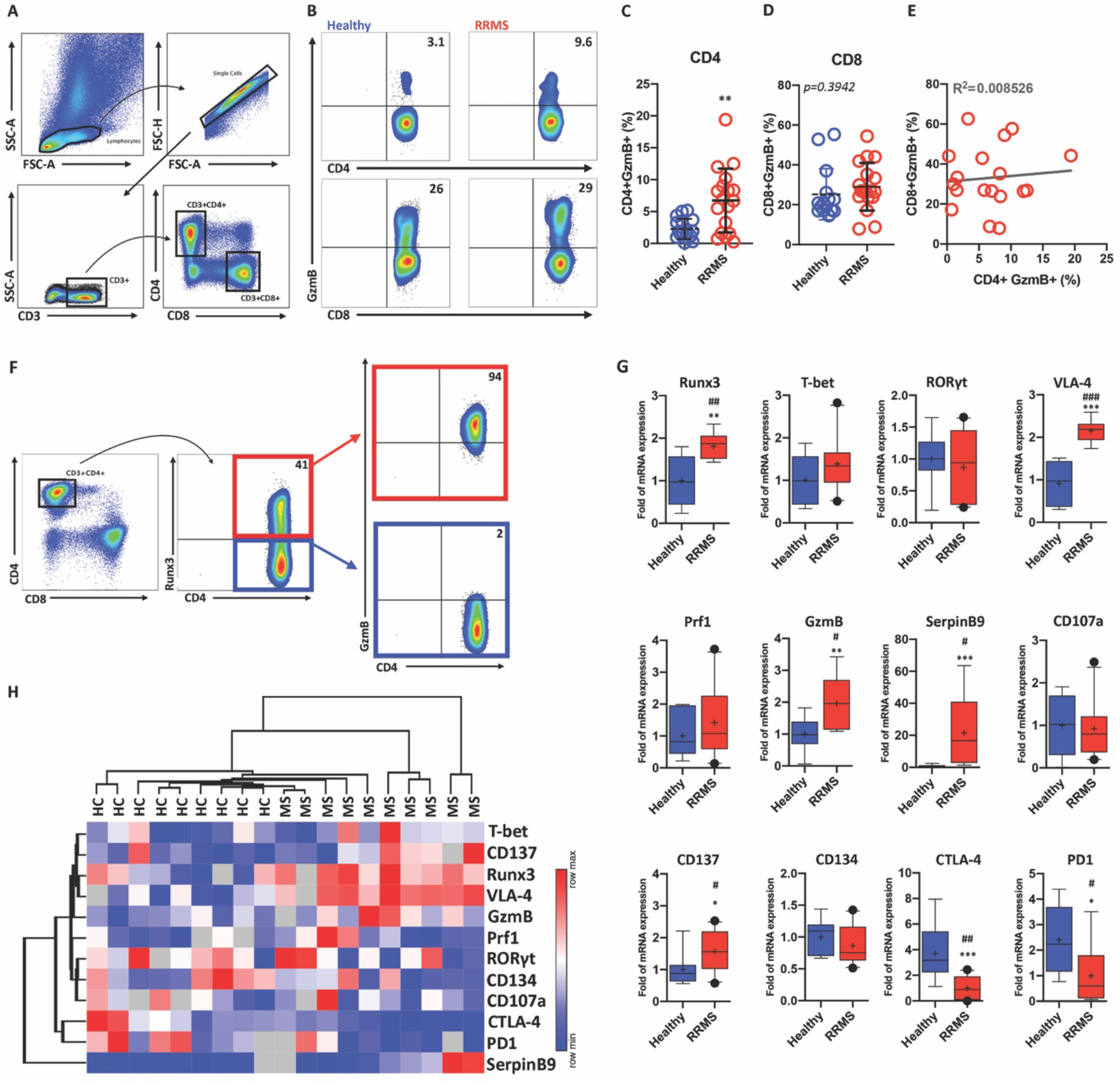
GzmB expression by CD4+ or CD8+ T cells in untreated patients with RRMS. (**A**) Gate strategy of flow cytometry analysis of CD3+CD4+ or CD3+CD8+ T cells from PBMCs from healthy individuals and RRMS patients. **(B)** Representative flow cytometry plot of GzmB expression in CD4+ or CD8+ T cells from PBMCs of healthy individuals and RRMS patients. **(C)** GzmB expression in gated CD3+CD4+ cells from PBMCs of HCs (blue) (*n=17*) or RRMS patients (red) (*n=18*). **(D)** GzmB expression in gated CD3+CD8+ cells from PBMCs of healthy individuals (blue) (*n=17*) or RRMS patients (red) (*n=18*). **(E)** Correlation of GzmB expression in CD3+CD4+ or CD3+CD8+ cells in RRMS (*n=18*). (**F**) GzmB expression in CD4+Runx3+ cells (red) or CD4+Runx3-cells (blue) (*representative from 5 patients*). (**G**) mRNA expression (Runx3, T-bet, RORγt, CD49d, SerpinB9, Prf1, CD107a and GzmB) of CD3+CD4+ sorted cells from PBMCs of healthy individuals (blue) (*n=9*) or RRMS patients (red) (*n=10*). (**H**) Hierarchical cluster analysis of the relative expression of Runx3, VLA-4, GzmB, Prf1, CD107a, SerpinB9, CD134, CD137, PD1, CTLA-4, T-bet and RORγt in sorted CD3+CD4+ T cells of PBMCs from healthy individuals (HC) (*n=9*) and RRMS patients (*n=9*). Data are represented in mean +/− SEM; *p<0.05, *p<0.05, **p<0.01, ***p<0.001; (Bootstrap resampling and permutation; #p<0.05, ##p<0.01, ###p<0.001).

Moreover, the presence of CD4+GzmB+ T cells is significantly higher in the PBMCs from RRMS patients assessed at the time of the diagnosis in comparison with those in the washout period (**Fig. S2A**). Although we cannot rule out the residual effect of the previous treatment in the washout group, this result indicates that the cytotoxic profile in CD4+ T cells is more prominent during the earlier phase of the disease. Indeed, secondary-progressive MS (SPMS) patients did not present increased levels of CD4+GzmB+ T cells in relation to HC (**Fig. S2B**). No significant difference was found for the CD8+ T cells population (**Fig. S2C and S2D**). Consistently with our data obtained in the experimental model, the expression of GzmB was almost restricted to Runx3-expressing CD4+ T cells (**Fig. 3F**).

Consistently, qPCR results demonstrated increased expression of Runx3 and GzmB mRNA in CD4+ T cells sorted from RRMS patients in comparison with HC (**Fig. 3G**). Although no significant difference was found in expression levels of T-bet or RORγt, there was a significant positive correlation (R^2^=0.6054) between mRNA expression of Runx3 and T-bet in RRMS patients (**Fig. S2E**). In contrast, the mRNA expression of Runx3 and RORγt trended (R^2^=0.3689; *p=0.1103*) to be negatively correlated (**Fig. S3E**). Taken together, these results reinforce the notion that CD4-CTL exhibits a Th1-like profile promoted by the mutual expression of Runx3 and T-bet (Cruz-Guilloty et al., 2009; Quezada et al., 2010; Serroukh et al., 2018; Y. Wang et al., 2014). Besides, we found an increased mRNA expression of SerpinB9, CD137, and VLA-4 in CD4+ T cells sorted from RRMS patients in comparison with HC (**Fig. 3G**). Interestingly, our results demonstrated a significant decrease in mRNA expression of CTLA-4 and PD1 in CD4+ T cells sorted cells from patients with RRMS, which agrees with previous work (Mohammadzadeh et al., 2018) (**Fig. 3G**). Together, these data indicate that, even in PBMCs, untreated RRMS patients present a specific enhancement of the cytotoxic profile in CD4+ T cells in comparison with HC. Indeed, hierarchal cluster analysis segregated RRMS patients and HCs based on the mRNA expression of cytotoxic-related and inflammatory molecules by CD4+ T cells (**Fig. 3H**).

### Cytotoxic Profile of Myelin-Reactive CD4+ T cells

Our data in the experimental model have indicated that the cytotoxic behavior of CD4+ T cells is enhanced upon autoantigen recognition (**Fig. 1A, 1B, and Fig. S1A**). Therefore, we accessed a publicly available sequencing (RNA-seq) expression dataset of myelin-reactive T cells from patients with MS (Cao et al., 2015). Briefly, CD4+CCR6+ memory T cells from HLA-DR4+ healthy individuals and MS patients were amplified and stimulated with myelin peptides (MOG_97-109_ and PLP_180-199_). After, myelin tetramers-positive (MOG_97-109_ and PLP_180-199_ tetramers) and tetramer-negative cells were sorted for RNA-seq (Cao et al., 2015). Comparison of tetramer-positive samples from HCs or MS revealed differential expression of transcription factors and activation markers previously related to CD4-CTL (Donnarumma et al., 2016; Śledzińska et al., 2020) or effector CD8+ T cells (D. Wang et al., 2018) (**Fig 4A**). Thus, we enriched the dataset analysis for cytotoxic-related and inflammatory molecules. Similarity matrix and correlation hierarchical clustering analysis show a strong correlation between MS tetramer-positive (TET+) samples based on the global expression of the selected genes (**Fig 4B**). Then, we used the similarity matrix of gene expression to select co-expressed genes (**Fig 4C**). Selected genes (46) were used to evaluate the expression in MS patients’ samples. Individually, 22 of 46 genes (45%) were significant between MS tetramer-positive (MS TET+) and MS tetramer-negative (MS TET-) samples (**Fig. 4D**). Nevertheless, based on these molecules, hierarchical clustering analysis segregate myelin-reactive MS TET+ cells from MS TET− (**Fig. 4D**). In order to have a better idea of the functional identity and connectivity of those molecules, we used the STRING network and clustering analysis (https://string-db.org/cgi/network.pl?taskId=rgnZ13we8jW). The molecules were grouped by molecular category (**Fig. 4E**). The analysis demonstrated a highly connected network, which is enriched by Th1 and Th17-related cytokines as well as cytotoxic-related effector molecules and transcription factors.

**Figure 4.**
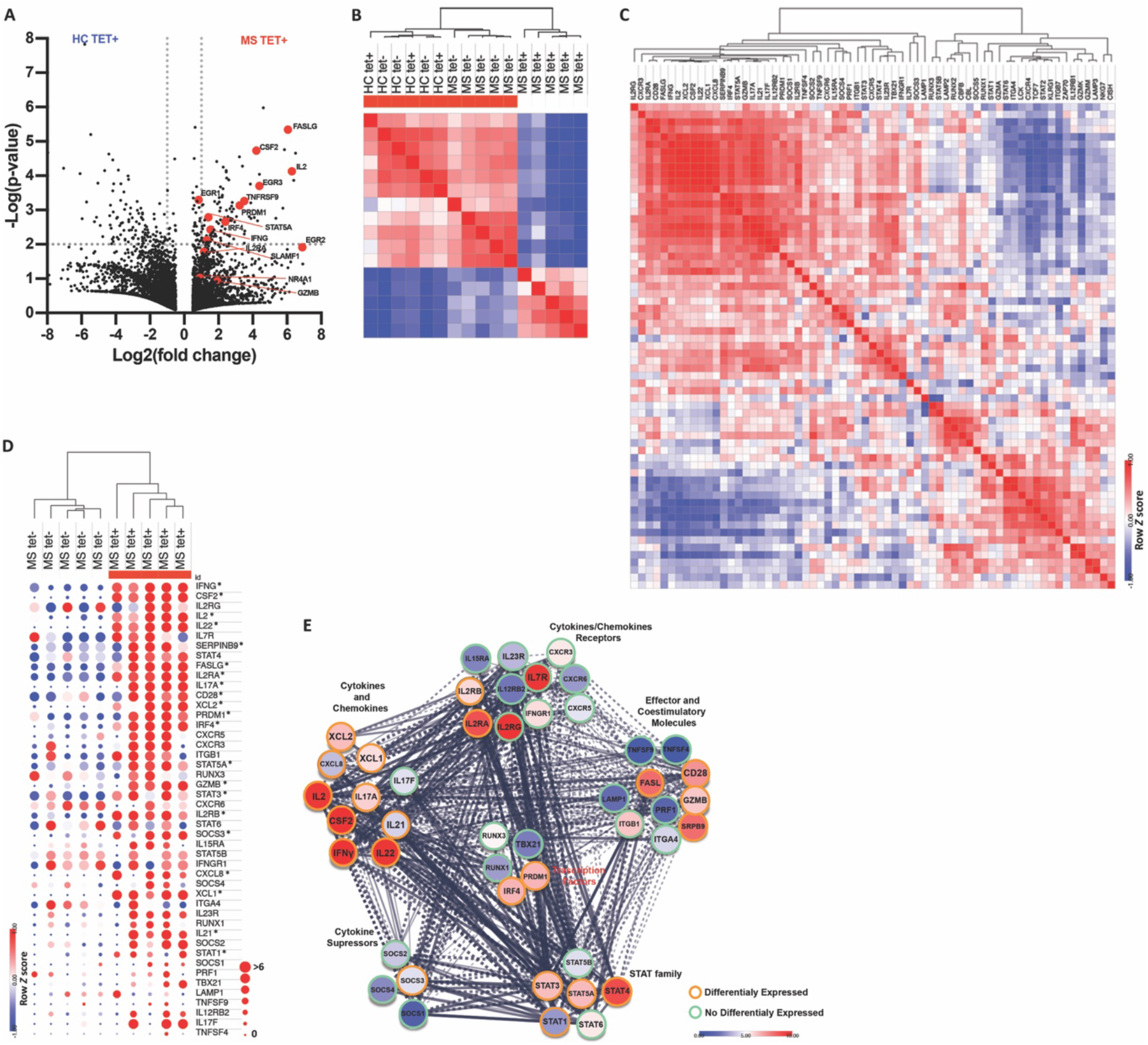
Cytotoxic profile of myelin-reactive CD4+ T cells. (**A**) Tetramer-positive cells differential gene expression between HC and RRMS samples (p < 0.01) (**B**) Similarity matrix and clustering analysis of global mRNA expression of CD4+CCR6+ memory T cells from MS tetramer+ (MS TET+), MS tetramer− (MS TET-), HC tetramer+ (HC TET+) or HC tetramer− (HC TET-) samples. (**C**) Similarity matrix and clustering analysis of selected genes expression in the 4 groups (**D**) Heat map of normalized log2FPKM values for the indicated genes in MS tetramer+ or MS tetramer− samples (z-score by color; log2 fold change by size - *differentially expressed). (**E**) STRING network representation of molecules enriched in MS tetramer+ samples. The color of each molecule shows a fold change of MS TET+ relative to MS TET-. Differentially expressed molecules are label by orange circles and no differentially expressed by green circles.

Altogether, our data consistently demonstrated a robust cytotoxic behavior of pathogenic CD4+ T cells upon autoantigen stimulus, which is restricted to patients with MS. Markedly, our analysis showed differential expression of IL2RA, PRDM1 (blimp1) and IRF4 (**Fig. 4E**), which are directly related to the effector activity of CD8+ T cells (D. Wang et al., 2018).

### Single-Cell mRNA Sequence of CSF Cells from MS Patients

Our data in the experimental model demonstrated that the cytotoxic profile of CD4+ T cells is enhanced after these cells reach the CNS. Therefore, we again took advantage of publicly available data to evaluate the extension of our data in CD4+ T cells from the CSF of MS patients. For that, we accessed the single-cell mRNA sequence (scRNA-seq) of CSF cells from MS-discordant monozygotic twin pairs (Beltrán et al., 2019a). First, we reconstructed the t-SNE leukocytes landscape (**Fig. 5A**). Interestingly, as in the original analysis, we were not able to segregate CD8+ and CD4+ T cells populations inside the T cell cluster. Instead, we also identify two regions, one housing preferentially CD4+ T cells and another housing CD8+ T cells (**Fig. 5B**). Therefore, we used the expression of CD3D and CD4 to manually select the CD4+ T cell population in order to evaluate the expression of cytotoxic-related molecules (**Fig. 5C**). Then, we used Louvain clustering to determine the subpopulations of CD4+ T cells (**Fig. S3A**). Further, we verified the expression of CD4-CTL-related molecules, defined by our data and others (Takeuchi et al., 2015; D. Wang et al., 2018), to identify which cluster represents better CD4-CTL population (**Fig. 5D**). The co-expression of those molecules indicates the cluster 3 (C3) is the CD4-CTL population (**Fig. 5E**). The expression of IL2RB, CRTAM, IFNG-AS1, EOMES, and GzmH were significantly higher in C3 in comparison with the remaining CD4+ T cells (**Fig. S3B**). C3 cluster presents a massive enrichment of CD4+ from patients with MS in comparison with HCs (Fig. 5F and S3C).

**Figure 5.**
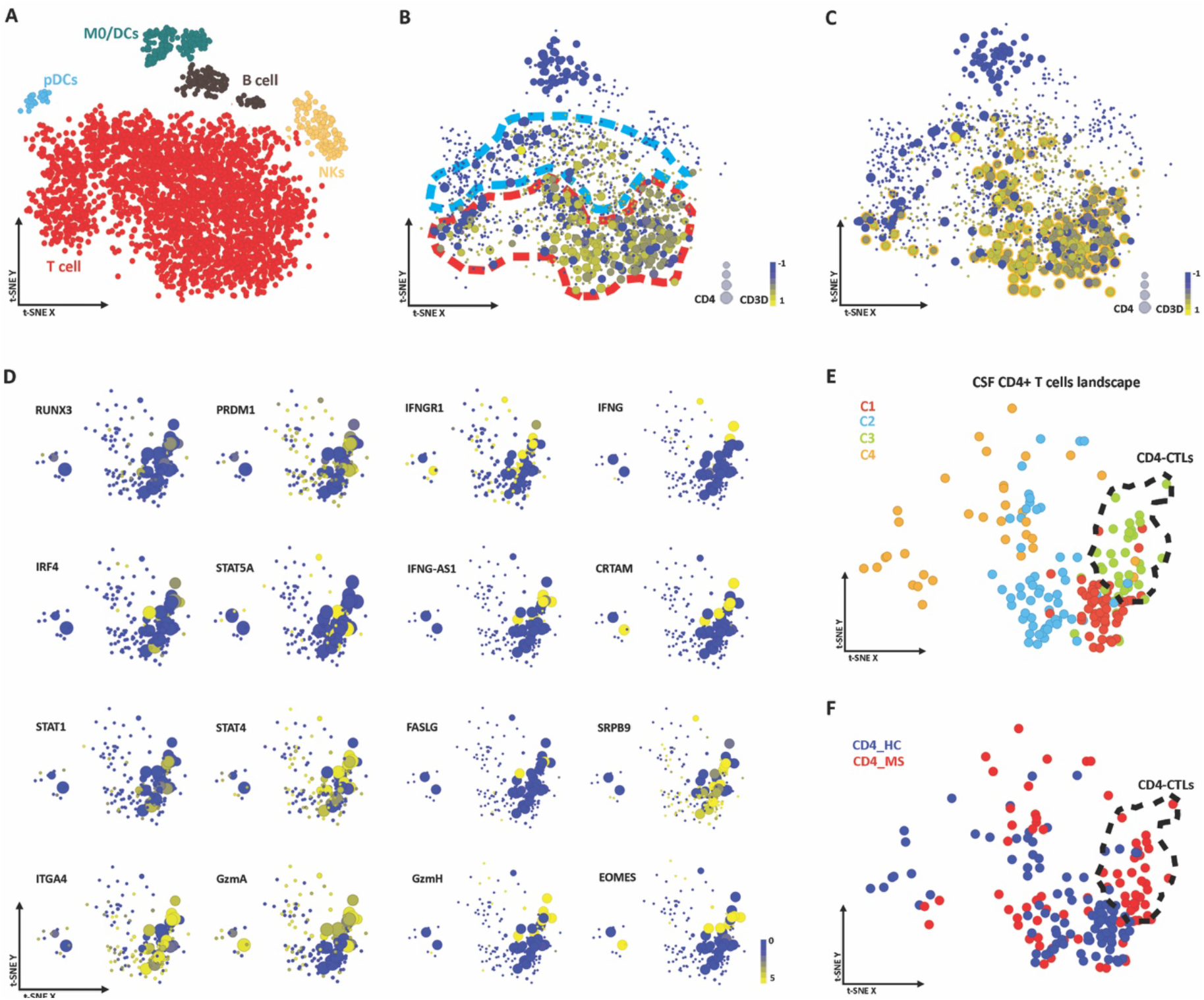
scRNA-seq of CSF cells. **(A)** Flowchart of the of t-SNE-based analysis of human CSF by scRNA-Seq. **(B)** Expression [log(Transcripts per million +1)] of CD3D (color) and CD4 (symbol size). The analysis shows two different populations in T cell cluster, one representing preferentially CD4+ T cells (red) and another representing preferentially CD8+ T cells (blue). (**C**) Selection (orange circle) of CD3D+CD4+ cells (CD3D by symbol color and CD4 by symbol size). (**D**) Expression of CD4-CTLs-related molecules in the CD3D+CD4+ population at the t-SNE-based landscape (IL2RB by symbol size; target gene by symbol color). (**E**) Louvain clustering of CD3D+CD4+ cells at the t-SNE-based landscape. Defined CD4-CTLs population (C3) were highlighted (discontinuous black line). (**F**) Distribution of CD4+ cells from CSF of HCs (blue) and RRMS patients (red) at CD3D+CD4+ t-SNE-based landscape.

Our analysis demonstrated that the CD4+ T cells of CSF from MS patients present an evident enrichment of CD4-CTLs in comparison with HC, again corroborating our data in the experimental model.

### Cytotoxic CD4+ T cells and VLA-4

Migration and BBB disruption are critical elements for the onset of autoimmune neuroinflammation (Kebir et al., 2007; Yednock et al., 1992). Collectively, our results in EAE or RRMS indicates a robust expression of VLA-4 by CD4-CTL. In monocyte-derived dendritic cells, the ectopic expression of Runx3 enhances the expression of VLA-4 (Domínguez-Soto et al., 2005). Consistently, we demonstrated a strong positive correlation (R^2^=0.4999) between the expression of Runx3 and VLA-4 in untreated RRMS patient CD4+ T cells (**Fig. S2E**). Moreover, ChiP-seq analysis from ENCODE data bank demonstrated direct binding of Runx3 to ITGA4 locus at the promoter region (**Fig. 6A**). In PBMCs from MS patients, the expression of GzmB is almost restricted to CD4+VLA-4+ cells (**Fig. 6B**). VLA-4 is the target of NTZ, which is a highly effective monoclonal antibody for the treatment of RRMS (Steinman et al., 2012). In this context, we hypothesized that NTZ-treatment might target specifically those CD4-CTLs. Therefore, to evaluate the impact of the treatments, specially NTZ, over CD4-CTL, we enrolled 48 treated RRMS patients (**Table 1**). Then, we evaluated the presence of CD4+GzmB+ and CD8+GzmB+ T cells in RRMS patients treated with immunomodulatory drugs [Glatiramer Acetate (GA) or interferon-β (IFN)], Fingolimod (FTY) or NTZ. As we hypothesized, NTZ-treated RRMS patients present enrichment of CD4+GzmB+ T cells in PBMCs in comparison with the other treatments or untreated patients (**Fig. 6C**). FTY-treated patients shown a slight increase in the percentage of CD4+GzmB+ T cells in comparison with IFN/GA-treated and untreated patients, which had been previously described (Fujii et al., 2016) (**Fig. 6C**). NTZ-treated patients also shown enrichment of CD8+GzmB+ T in comparison with the other treatments or untreated patients (**Fig. 6D**).

**Figure 6.**
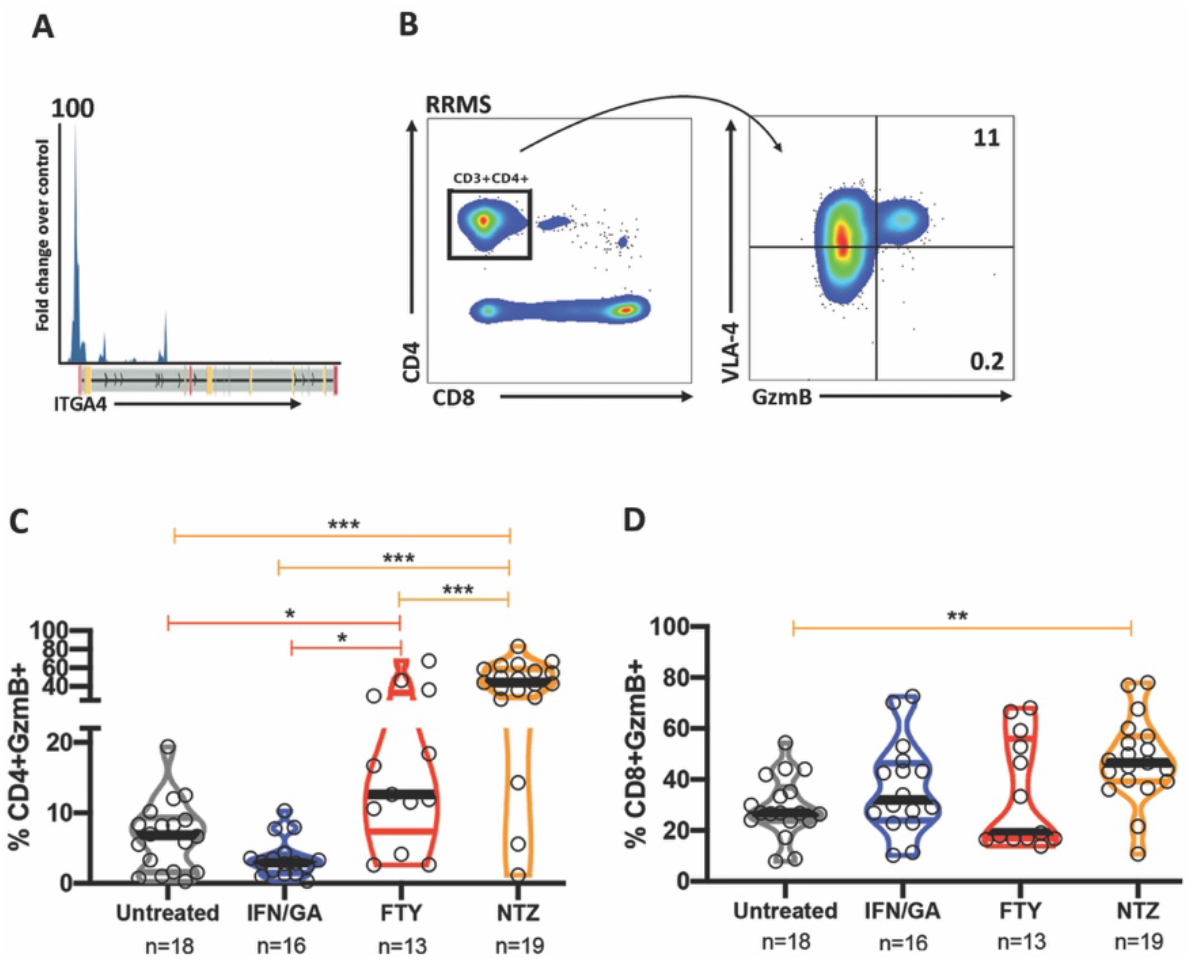
Cytotoxic CD4+ T cells and VLA-4. (**A**) Runx3 ChiP-seq analysis from ENCODE data bank in ITGA4 locus. (**B**) VLA-4 and GzmB expression (right) in gated CD3+CD4+ (left) from PBMCs of patients with MS (*representative from 5 patients*). (**C**) Violin plot of GzmB expression in gated CD3+CD4+ cells from PBMCs of untreated (gray) (*n=18*), IFN-treated (blue) (*n=16*), FTY-treated (red) (*n=13*) and NTZ-treated (orange) (*n=19*) RRMS patients. (**D**) Violin plot of GzmB expression in gated CD3+CD8+ cells from PBMCs of untreated (gray) (*n=18*), IFN-treated (blue) (*n=16*), FTY-treated (red) (*n=13*) and NTZ-treated (orange) (*n=19*) RRMS patients. Data are represented in median (bold bar) and 95% C.I. (color bars); *p<0.05, **p<0.01 ***p<0.001.

## Discussion

Here, we demonstrated that autoaggressive encephalitogenic CD4+ T cells present a cytotoxic phenotype during the early phase of EAE and RRMS. Robust and concordant data support the critical role of CD4-CTLs during initial autoimmune neuroinflammation.

Our mRNA expression analysis of CD4+ T cells from EAE or MS patients demonstrated an enhancement of cytotoxic-related molecules. Also, encephalitogenic CD4+ T cells were sufficient to promote the initial CNS damage, in the absence of a previous inflammatory milieu. Of note, CD4+ T cells from PBMCs of patients with MS show cytolytic activity to neurons and oligodendrocytes *in vitro* (Kebir et al., 2007; Zaguia et al., 2013). Therefore, CD4+ T cells present not only a cytotoxic phenotype but also a direct cytolytic activity.

Previously, autoaggressive Th17 cells have been shown to use GzmB to breach the BBB during EAE initial neuroinflammation (Kebir et al., 2007). In contrast, our data demonstrated that CD4-CTLs presents a Th1 profile preferentially. However, in the EAE model, Th17 cells turn into IFNγ-producing cells after reach the CNS (Hirota et al., 2011). Thus, effector CD4-CTLs inside the CNS and autoaggressive Th17 cells in the periphery may represent the same population in different stages. Consistently, the expression analysis of myelin-reactive CD4+ T cells from patients with MS showed an enhancement of Th1 and Th17-related cytokines as well as cytotoxic-related molecules. Moreover, our flow cytometry analysis of CNS-infiltrated CD4+ T cells presents a population that expresses high levels of VLA-4 and intermediary levels of GzmB (VLA-4^hi^). VLA-4 is an adhesion receptor essential to the transmigration of circulating leukocytes into the CNS in EAE (Yednock et al., 1992). Thus, the VLA-4^hi^ population is probably the CD4+ T cells that just infiltrated the CNS. In this case, the differential expression of GzmB by this population is in agreement with Kebir’s data (Kebir et al., 2007).

The conversion of Th17 cells into IFNγ-producing cells is dependent on T-bet and Runx transcription factors expression (Y. Wang et al., 2014). In this context, our data revealed that the expression of GzmB by CD4+ T cells is almost restricted to Runx3-expressing cells either in EAE or RRMS. Noteworthy, the heterogeneous network edge prediction analysis indicates Runx3 as an MS-associated gene (Himmelstein and Baranzini, 2015). Upon TCR stimulation, Runx3 promotes accessibility to regions highly enriched with IRF and PRDM1-like motifs in naïve CD8+ T cells (D. Wang et al., 2018). Subsequently, the expression of those transcription factors and IL-2 receptor is critical to early effector and memory precursor CD8+ T cell differentiation (D. Wang et al., 2018). Consistently, our RNA-seq analysis demonstrated an enhancement of PRDM-1, IRF4, IL2RA, and IL2RB expression either by myelin-reactive T cells or CD4-CTLs from CSF. Remarkable, the ablation of PRDM-1 prevent GzmB expression and anti-tumor activity of CD4-CTLs (Śledzińska et al., 2020). Moreover, IRF4 and PRDM-1 deficient mice present a minor incidence and severity of EAE, very similar to what we have shown to Runx3 deficient mice (Brüstle et al., 2007; Jain et al., 2016). Besides the enhancement of cytotoxic-related transcription factors, we found a differential expression of cytotoxic effector molecules such as; GzmB, GzmA, SerpinB9, CD137 (TNFRSF9), and FASLG either in peripheral blood or CSF. Altogether, our data robustly demonstrated that autoaggressive CD4-CTLs fulfills the majority of the molecular pathways of effector CD8+ T cells (Hartung et al., 2015; Janas et al., 2005; Pearce et al., 2003; Suzuki et al., 1997; Verdeil et al., 2006; D. Wang et al., 2018; Zhang et al., 2019).

Strikingly, we and others were unable to distinctly separate CD4+ and CD8+ T cell populations from MS samples by their expression profile (Beltrán et al., 2019b; Schafflick et al., 2020). These data contribute to the notion that CD4+ T cells behavior much like effector CD8+ T cells during the course of the disease. In MS, the percentage of cytotoxic-like CD4+ T cells (CD4+CD28null) in the peripheral blood seems to present a direct link with the disease severity as well as a prognostic value (Peeters et al., 2017). Consistently, we demonstrated that the presence of CD4+GzmB+ T cells is prominent in the peripheral blood of untreated patients with RRMS assessed at the very early stage of the diagnosis. Moreover, the expression of cytotoxic-related molecules by CD4+ T cells was sufficient to segregate untreated RRMS and HC. Thus, although our data need to be extended to a larger cohort, the cytotoxic profile of CD4+ T cells holds great potential to aid RRMS diagnosis.

Further, we found that NTZ treatment massively restrains CD4+GzmB+ and CD8+GzmB+ T cells in the peripheral blood. The discontinuation of NTZ treatment may lead to a clinical and radiological rebound, which, in some cases, is fatal (Larochelle et al., 2017). In fatal rebound following NTZ discontinuation CD4+GzmB+ and CD8+GzmB+ T cells accumulate in the brain parenchyma (Larochelle et al., 2017). Noteworthy, the high cytotoxic activity of lymphocytes has been related to steroid resistance in obstructive pulmonary disease and neuromyelitis optica spectrum disorder (Boldrini et al., 2020; G. Hodge and S. Hodge, 2016). Thus, the restraining of those cytotoxic cells in the periphery seems to be, at least in part, direct related to the beneficial effect of NTZ treatment. In this context, the percentage of CD4-CTLs in the peripheral blood could reveal a valuable readout for monitoring the treatment outcome.

Our finds suggest a critical role of CD4-CTLs during the initial autoimmune neuroinflammation. Furthermore, our results argue for the diagnostic and prognostic potential of monitoring CD4-CTLs percentage in the peripheral blood. A comprehensive assessment of the cellular and molecular mechanisms of CD4-CTLs might be beneficial to understand their pathological role in MS better. Ambitiously, further studies might point to CD4-CTLs as a viable target for the development of new therapeutic strategies.

## Materials and Methods

### Study approval

All animal experiments in this study followed protocols approved by the Animal Care and Use Committee at the University of Campinas (CEUA-UNICAMP #4657-1, #4656-1, #4658-1, #3890-1, #4420-1, #3936-1 and GMO CiBio #01/2013). Both RRMS patients and healthy control subjects included in this work signed a term of consent approved by the University of Campinas Committee for Ethical Research (CAAE: 53022516.3.0000.5404 and CAAE: 64431516.4.0000.5404).

### Animals

Eight-week-old C57BL/6 background mice (C57BL/6^WT^, IFNγ^*yfp*^(*B6.129-Ifng^tm3.1Lky^/J*), IL-17A^*gfp*^(*C57BL/6-Il17a^tm1Bcgen^/J*), Runx3^fl/fl^(*B6.129P2-Runx3^tm1Itan^/J*), Runx3^*yfp*^(*B6;129P2-Runx3^tm1Litt^/J*), and CD4^Cre^(*STOCK Tg(Cd4-cre)1Cwi/BfluJ*)) were obtained from the Jackson Laboratory and established as a colony at the University of Campinas Breeding Center, where they were housed and maintained under pathogen-free conditions in the university animal facility. The experimental animals were allowed access to standard rodent chow and water ad libitum, with the temperature maintained between 21° and 23°C and a 12-h light/12-h dark cycle. All procedures were carried out following the guidelines proposed by the Brazilian Council on Animal Care.

### Subjects

Untreated RRMS patients were recruited at the time of the diagnostic (8) or during treatment washout (10). Treatment washout was applied due to therapeutic failure or after 24 months of Natalizumab treatment (Stangel and Stüve, 2014) (**Table S1**). Also, 48 treated (16 IFN/GA, 13 FTY, and 19 NTZ) RRMS patients were recruited (**Table S1**). All RRMS patients were included in the study after confirmation of the diagnosis according to the revised McDonald criteria (Polman et al., 2011). Also, 17 matched healthy subjects were included in the control groups.

### Blood samples collection and lymphocyte separation

Peripheral blood (25 mL) samples were collected from RRMS patients and healthy volunteers. Peripheral blood mononuclear cells (PBMCs) were separated by Ficoll-Hypaque^®^ density gradient centrifugation, as previously described (Longhini et al., 2011).

### Experimental Autoimmune Encephalomyelitis

Mice were immunized subcutaneously with 150 μg of MOG_35-55_ or 100 μg of OVA in complete Freud’s adjuvant (CFA), and 4 mg/ml heat-inactivated *Mycobacterium tuberculosis* and with an intraperitoneal injection of Bordatella pertussis toxin (0 and 2 d.a.i.).

### Quantitative PCR

mRNA was extracted using the RNeasy micro kit (QIAGEN) and reverse transcribed to cDNA. For mouse experiments, the primers for RORα, RORγt, T-bet, Eomes, Runx3, IFNγ, IL-17a, GzmB, GzmC, Xcl-1, CD134, CD137, CD107a, CD107b, PRF1, and VAV3 were obtained from Applied Bioscience inventoried catalog. For human samples, qPCR was performed using SYBR^®^ Green manufacturer’s instructions (BioRad, USA). PCR analysis was performed using a TaqMan ABI Prism 7500 Sequence Detector (PE Applied Biosystems, Germany), and mRNA was normalized to that of a housekeeping gene (GAPDH or HRP). The data were generated using independent duplicate measurements. The threshold cycle value of individual measurements did not exceed 0.5 amplification cycles.

### Antibodies and Flow Cytometry

The following antibodies were used: for mouse; anti-CD3 (145-2C11), anti-TCRαβ (H57-597), anti-CD4(GK1.5), anti-CD8α (53-6.7), anti-GzmB (NGZB), anti-CD134 (OX-86), anti-CD107a (1D4B), anti-Eomes (Dan11mag), anti-Runx3 (R3-5G4), anti-VLA-4 (R1-2) and anti-ThPOK (2POK); for humans; anti-CD3 (SK7, SP34-2), anti-CD4 (SK3, RPA-T4), anti-CD8 (SK1), anti-Runx3 (R3-5G4), anti-VLA-4 (9F10) and anti-GzmB (GB11). Antibodies were conjugated to the following fluorescent dyes: fluorescein isothiocyanate (FITC), phycoerythrin (PE), peridinin chlorophyll protein (PercP), allophycocyanin (APC), phycoerythrin-cyanine 7 (PECy7), allophycocyanin-cyanine 7 (APCcy7), Brilhant-Violet 421 (BV421). A Foxp3 transcription factor fixation/permeabilization kit (eBioscience, USA) was used for intracellular staining of Eomes, Runx3, CD107a, ThPOK, and GzmB. All analyses were performed using a flow cytometer Gallios (Beckman-Coulter, USA) or FACSVerse (BD bioscience, USA).

### Cell sorting

All sorting was performed using a cell sorter flow cytometer FACSAria II (BD Bioscience, USA). Cells were kept on ice before and after sorting analysis. Cell purity was confirmed immediately after sorting.

### Brain slice incubation

Brains were aseptically obtained, rapidly packed in 2% agarose cubes and sliced (400 μm) with a vibratome. CD4+ T cells from mice immunized for EAE or controls (ovalbumin immunized mice or unimmunized mice) were sorted with a flow cytometer. Sorted cells were incubated with slices (5×10^5^ cells/slice) for 6 hours in Hanks’ Balanced Salt solution (Sigma-Aldrich, USA) supplemented with 1% of penicillin/streptomycin solution (Thermo-Fisher Scientific, USA). After the incubation period, brain slices were placed in Tissue-Tek OCT compound (Sakura Finetek, USA) and frozen.

### Confocal Microscopy

*In situ* expression of cleaved caspase-3 in the brain from EAE and from control animals (ovalbumin immunized mice or unimmunized mice) was performed following the protocol for immunofluorescence staining. Briefly, frozen brains in Tissue-Tek OCT compound (Sakura Finetek, USA) were sliced to produce 10 μm sections using a cryostat and submitted to staining. For this, the tissue was blocked to avoid nonspecific reactions with donkey serum (Sigma-Aldrich, USA) and permeabilized with Triton X-100 (Sigma-Aldrich, USA). Then, a primary antibody against cleaved caspase-3 (Cell Signaling Technology, USA) was added (1:200) and incubated overnight. The following day, slices were washed and incubated with secondary antibodies (Alexa 488) (Cell Signaling Technology, USA) (dilution 1:500) plus DAPI (Sigma-Aldrich, USA). For confocal controls, primary antibodies were omitted from the staining procedure and were negative for any reactivity. All images were obtained using confocal microscopy at INFABIC-UNICAMP.

### RNA-seq data analysis

We accessed publicly RNA-Seq expression data deposited in the NCBI’s Gene Expression Omnibus (GEO) database under accession numbers GSE66763. Briefly, CCR6+ memory CD4+ T cells from HLA-DR4+ healthy controls (HC) and HLA-DR4+ MS patients (MS) were amplified by PHA and IL-2 and stimulated by irradiated autologous monocytes and DR4 myelin peptides (MOG97–109 and PLP180–199). Then, myelin tetramer+ (TET+) and tetramer− (TET-) cells were sorted for RNA sequencing (Cao et al., 2015).

### scRNA-Seq data analysis

We accessed publicly scRNA-Seq expression data deposited in the NCBI’s Gene Expression Omnibus (GEO) database under accession numbers GSE127969. Then, we used the same filter parameters (removing transcripts detected in fewer than 3 cells and kept only cells with 200 to 6000 detected transcripts) from the original publication (Beltrán et al., 2019a). The detection of variable genes, unbiased graph-based clustering of single-cell data, was done using scOrange software (Stražar et al., 2019).

### Statistical analysis

Statistical significance of the results was determined using analysis of variance (Kruskal–Wallis or ANOVA; U-test or *t*-test). Grubbs’ test was used to determine and exclude outlier values. Correlations were determined by Pearson or Spearman rank tests as appropriate. Bootstrap resampling (100,000x) and permutation (BsRP) were used in the qPCR analysis of human samples to minimize statistical interference due to the small sample size. The results of BsRP have expressed *p*-value (n*H*_0_+1/99999+1). Hierarchical cluster analysis was performed using the Euclidean distance metric. A P-value of less than 0.05 was considered significant in all tests. t-distributed stochastic neighbor embedding (t-SNE) (iteration = 3,000; perplexity = 30; learning rate = 252; gradient algorithm = Barnes-Hut) was performed using Flowjo R studio-based plugin.

### Software

Basic statistical analysis was performed using GraphPad Prism software. Bootstrap resampling and permutation were performed using Rstudio software. Heat map and hierarchical cluster analysis were performed using the MORPHEUS web-based tool (https://software.broadinstitute.org/morpheus/). All images obtained were analyzed using the Image J. FACS data was analyzed using the FlowJo 10.6 software package (Tri-Star, USA). Single-cell data analysis was done using scOrange 3.24 software.

## Data Availability

All raw data are publicly available at the University database (https://drive.google.com/open?id=1-0PoClxU8m4hjW75REwIv0Kn7WbxkANs&authuser=neuroib@unicamp.br&usp=drive_fs).

## Acknowledgment

The authors would like to thank M.A. Mori, N.O.S. Câmara, D. Martins-de-Souza, H.M. Souza, P.M.M. Vieira, M.A. Vinolo, F. Papes, R. Vicentini and H.F. Carvalho for critically reading the manuscript and/or for providing resources. The authors acknowledge the aid of Natalia Coutouné with bootstrap resampling and permutation script design. We thank the National Institute of Science and Technology of Photonics Applied to Cell Biology (INFABIC) for aid with confocal microscopy. The authors acknowledge the technical help of Adriel S. Moraes, Marcos C. Menegheti.

## Author Contributions

F.P., A.M.M., and G.A.D.M. performed most of the experiments in the experimental model. VOB performed most of the experiments in RRMS patients and healthy donors. C.R.V.S, F.vG., and A.D. selected and recruited all patients and healthy individuals. C.F.R. performed acute slice incubation and immunohistochemistry experiments. E.S.M.F., R.S.C., N.B., B.B.C., and V.C.L. established and performed experiments with genetically modified mouse strains. M.B-jr. performed experiments with OVA antigen-immunized animals. A.L.F.L. and M.R-P. designed and performed flow cytometry and cell sorting experiments. A.S.F. and L.M.B.S designed the experimental work. A.S.F. coordinated the study and wrote the manuscript with inputs from co-authors.

## Supplementary Material

**Figure S1.**
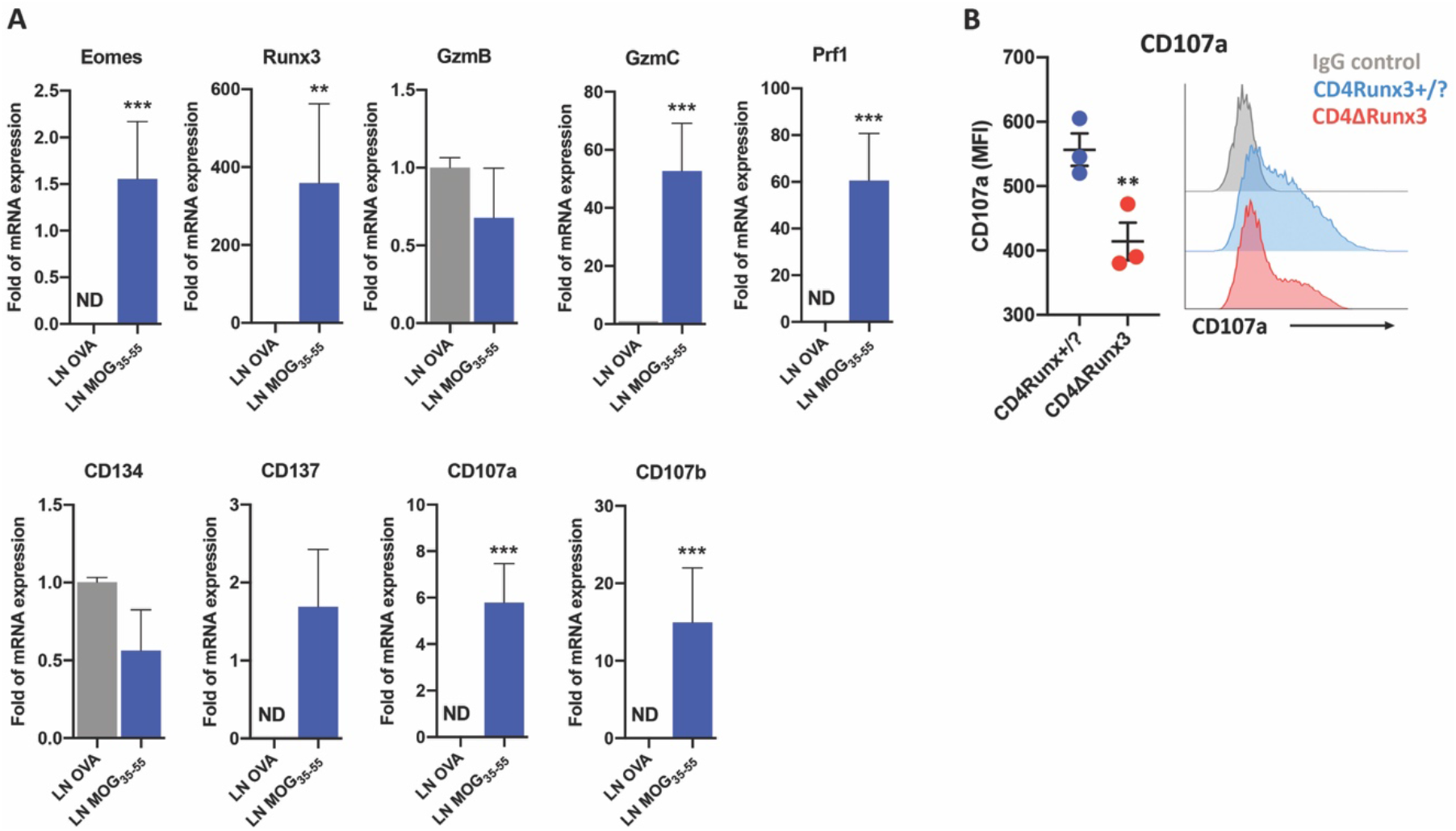
Expression of cytotoxic-related molecules in CD4+ T cells from lymph nodes. (**A**) mRNA expression of cytotoxic-related molecules in CD4+ cells sorted from the lymph nodes of OVA-immunized mice 10 d.a.i. (blue), or lymph nodes of MOG_35-55_-Immunized mice; 10 d.a.i. (red) (*n=3-9*). (**B**) CD107a expression (MFI) in CD3+CD4+ cells from CD4Runx3+/? (blue) or CD4∆Runx3 mice 10 d.a.i. (red). Data are represented in mean +/− SEM; *p<0.05, **p<0.01, ***p<0.001.

**Figure S2.**
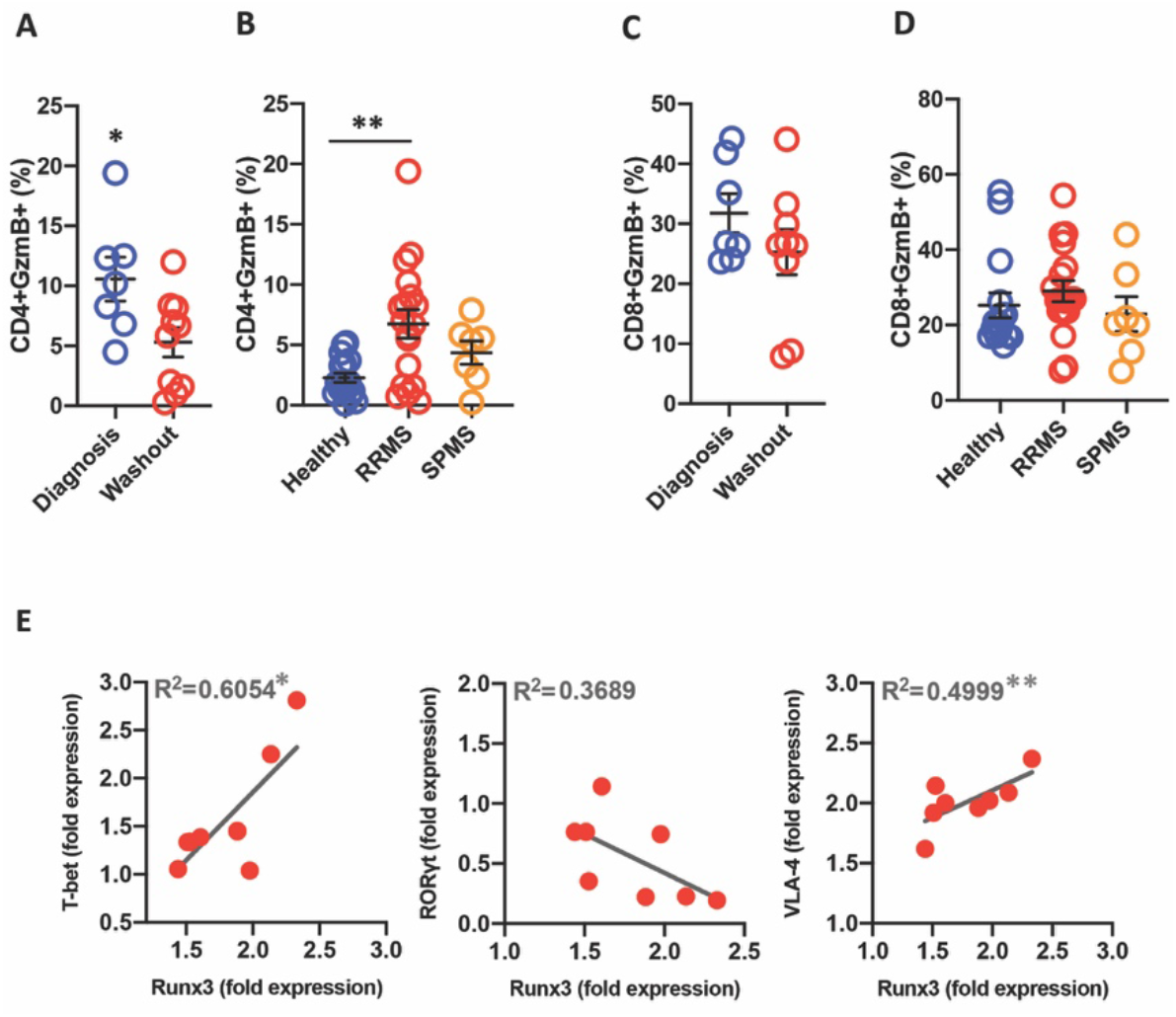
GzmB expression by CD4+ or CD8+ T cells RRMS or SPMS patients. (**A**) GzmB expression in gated CD3+CD4+ cells from PBMCs of untreated RRMS assessed at the time of diagnosis (blue) (*n=8*) or untreated RRMS patients assessed at washout period (red) (*n=10*). (**B**) GzmB expression in gated CD3+CD4+ cells from PBMCs of HCs (blue) (*n=17*), RRMS patients (red) (*n=18*) or SPMS patients (orange) (*n=7*). **(C)** GzmB expression in gated CD3+CD8+ cells from PBMCs of untreated RRMS assessed at the time of diagnosis (blue) (*n=8*) or untreated RRMS patients assessed at washout period (red) (*n=10*). (**D**) GzmB expression in gated CD3+CD8+ cells from PBMCs of HCs (blue) (*n=17*), RRMS patients (red) (*n=18*) or SPMS patients (orange) (*n=7*). (**F**) Correlation analysis of mRNA expression of T-bet, RORγt or VLA-4 with Runx3 in CD3+CD4+ T cells sorted from PBMCs of RRMS patients. Data are represented in mean +/− SEM; *p<0.05, **p<0.01

**Figure S3.**
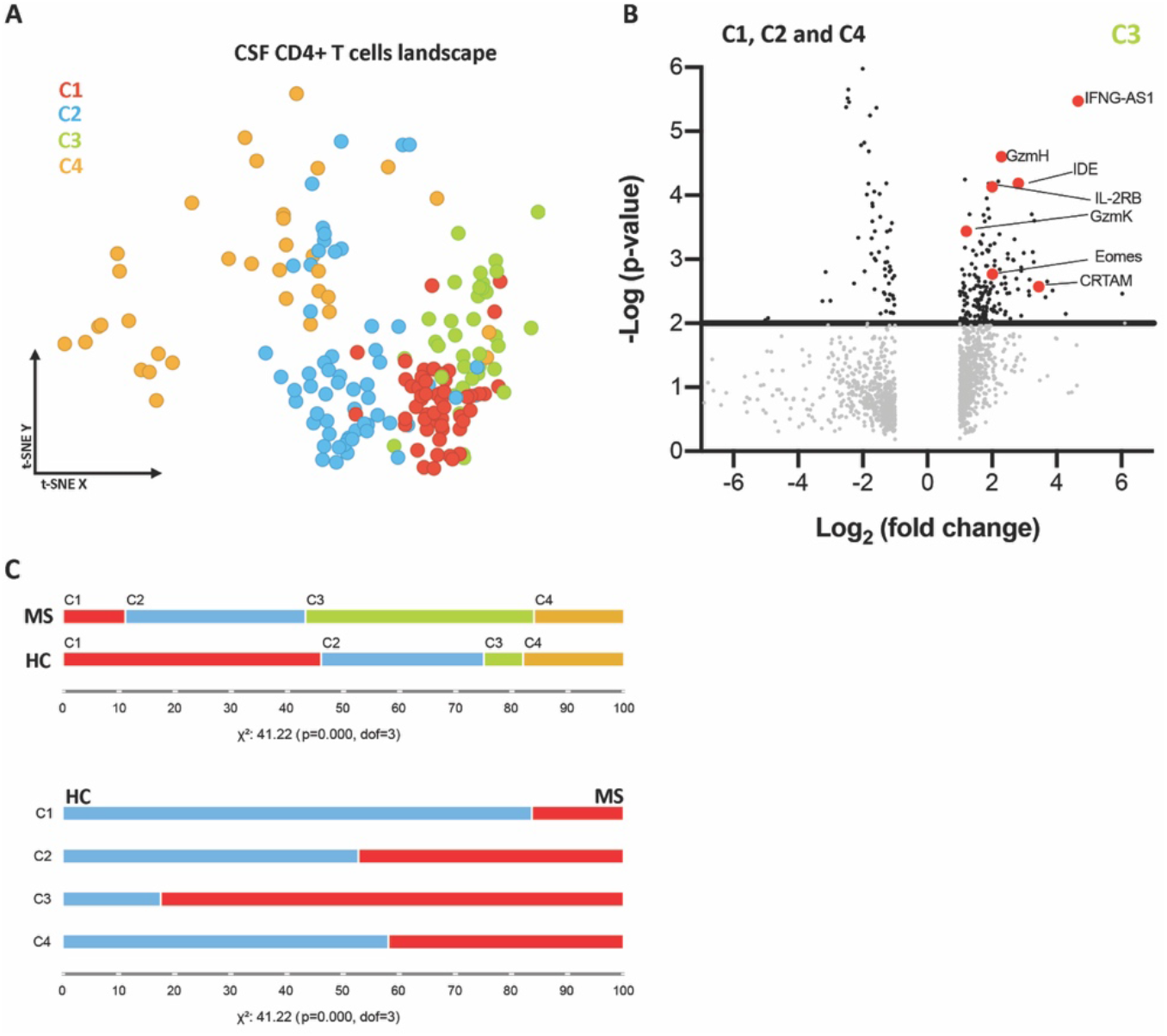
GzmB expression by CD4+ or CD8+ T cells RRMS or SPMS patients. (**A**) Louvain clustering of CD3D+CD4+ subpopulations at the t-SNE-based landscape. (**B**) Volcano plot of global mRNA expression of C3 cluster cell in relation to remaining cells (C1, C2, and C4). (**C**) Frequency of each Louvain cluster in CD3D+CD4+ cells from MS patients or HCs (top). Frequency of CD3D+CD4+ cells from MS patients or HCs in each Louvain cluster (bottom).

